# A zebrafish model of developmental joint dysplasia: Manipulating the larval mechanical environment to drive the malformation and recovery of joint shape

**DOI:** 10.1101/155911

**Authors:** Karen A Roddy, Roddy EH Skinner, Lucy H Brunt, Erika Kague, Stephen Cross, Emily J Rayfield, Chrissy L Hammond

**Affiliations:** School of Physiology, Pharmacology and Neuroscience, University of Bristol, BS8 1TD, U.K.; School of Earth Sciences, University of Bristol, BS8 1TD, U.K.

## Abstract

Developmental dysplasia of the hip (DDH), a malformation of the acetabulum, is a frequent cause of early onset osteoarthritis. The disease encompasses a spectrum of severities, some of which are more amenable to treatment. Embryonic immobilisation significantly impairs the development of joint shape however the impact of this malformation to the function and growth of the joint in the short to medium term is unclear. We developed a novel model of developmental joint dysplasia using the zebrafish jaw joint to identify the mechanisms regulating cellular plasticity and ability to recover joint shape and function. Larval zebrafish were immobilised either pharmacologically or using targeted ablation of jaw muscles to induce an altered joint shape. Following restoration of muscle activity we dynamically monitored the joint shape and function in individuals at cellular resolution impossible in other vertebrate species. Reflecting the variability of the human condition we found a proportion of joints will recover both their shape and function, while others will not; despite coming from a genetically homogenous population. This allowed us to study what controls likelihood of recovery; we identified a number of cellular changes that predict likelihood of functional recovery, including position of precursor cells, and specific patterns of proliferation, migration and differentiation in joints and associated connective tissues. These factors together predict recovery better than severity of malformation alone. Using Finite Element Analysis we studied the mechanics of joints representative of ones that recover and those that fail to identify differences in patterns of strain that could explain the cellular behaviours that underpin likelihood of recovery. Thus, this model would enable the study of the short to long term impact of altered joint shape on function and could help to identify the changes that render an individual more receptive to treatment and therefore may potentially be indicative of long term joint health.

## Introduction

The complex shape of a synovial joint is a vital component of its function; governing its range of motion and capacity to efficiently transmit load. Congenital joint shape defects, such as developmental dysplasia of the hip (DDH) increase an individual’s risk of osteoarthritis (OA) even if treated successfully; although some subtle forms of DDH can resolve spontaneously [1]. DDH, caused by a shallow acetabulum leading to a joint prone to subluxation is associated with factors that constrain foetal and postnatal movement including breech presentation and swaddling [2]. DDH has a dramatic impact on the hip joint but more subtle variations in joint shape have been associated with increased risk of developing osteoarthritis [3,4]. Thus, joint shape is an important regulator of joint fitness. Current treatment for DDH using a Pavlic harness relies on postnatal remodelling and growth to modify the shape of growing joint, however, success is linked to early treatment and failure to surgical intervention [1]. The mechanisms through which these abnormal joint shapes are corrected, and the factors governing the optimal window for treatment remain unclear.

Mechanoregulatory signals generated by muscle contraction are vital to the development of its characteristic 3D structure [5–9] and have been suggested as primary driver of DDH development [10]. Studies in the chick [6,11,12], mouse [13–15] and zebrafish [5,15,16] have shown that loss of muscle activity leads to malformed joints lacking features such as ligaments, articular cartilage, meniscus and articular processes. Gradients of shear, tension and compression are created during muscle contractions and cells are differentially responsive to these varying mechanical signals altering their cell behaviour or differentiation in response to the changing mechanical force [17–19]. Mechanical stimulus is also a major modifier of disease progression in OA. Immobilising murine knee joints after surgical destabilisation, reduces expression of OA associated genes and protects these joints from degeneration [20] and also slows tissue damage in patients with established OA [21,22]. However, immobilisation can also drive the degeneration of healthy cartilage [23]. Thus, mechanical stimulus wrongly applied can be detrimental to joint health. How aberrant mechanical stimuli, generated by the movement of a malformed joint might be integrated into the regulatory framework governing postnatal growth in order to achieve joint recovery is poorly understood.

Various models of DDH exist; including spontaneous canine models [24,25], cast/swaddling models in rats and rabbits [26,27] and foetal immobilisation [10]. Canine and cast/swaddling models do not recapitulate the initial development of DDH as the hip is normal at birth. The chick immobilisation model can induce a shallow acetabulum however joint degeneration has not been confirmed. Here we present a zebrafish model that facilitates the study of the early developmental steps leading to a subluxed joint and the testing of factors underpinning likelihood of recovery. Zebrafish develop externally making them tractable for drug treatment or laser surgery. Zebrafish have synovial joints and have been demonstrated to develop phenotypes resembling osteoarthritis as they age [28,29]. Most importantly the availability of transgenic lines labelling the developing skeleton enables studies of cell behaviour in live larvae [39}. We hypothesise that restoration of movement, following immobilisation, drives morphogenetic changes in the zebrafish jaw and joint remodelling. To test this, we induced jaw joint malformation using a period of pharmacological immobilisation [5] or by muscle ablation, then allowed movement to resume and tracked the changing jaw shape, cell behaviour and differentiation. Laser ablation has been used to study repair mechanisms after injury [30] and was used here to sever the adductor mandibularis. We show that immobilised larvae that recover have a significantly different joint shape to those that fail to recover; and they successfully remodel the joint and associated tissues including the ligament and enthesis. We also demonstrate that recovery potential is underpinned, in part, by changes to cell orientation, migration and proliferation. Using finite element analysis to explore the mechanoregulatory environment experienced by recovering and subluxed larvae we identify <u>differences.in</u> patterns of strain that could explain the cellular behaviours that underpin likelihood of recovery.

## Results

### Resumption of movement following immobilisation leads to changes in jaw placement

The anaesthetic MS222 (tricaine) induces flaccid paralysis in zebrafish and has been shown to lead to malformation of the jaw joint and changes to joint cell behaviour [5,15,16]. To determine the capacity of these malformed joints to recover functionality larvae were treated with MS222 for 48hrs from 3-5dpf then allowed to resume movement. To rule out secondary drug effects and to investigate the impact of targeted change to jaw mechanics we also developed a model in which we bilaterally ablated the adductor mandibularis, which is responsible for jaw closure, using a laser. Following ablation, muscle regrowth was visible by 1 day post ablation (dpa) and muscle fibres reconnected the elements and were contractile by 2dpa (at 5dpf) (S1 Fig).

Following either anaesthesia or muscle ablation the jaw joint was subluxed in all larvae such that it was rendered unable to close (S1-3 Video) at 5dpf (Fig 1B-C,1-2). Underlying this loss of function was a significant change to the shape of the MC and jaw joint which lacked the complementary surfaces of control joints (Fig 1A-C, 3-4). Rather, the MC overlapped the palatoquadrate on the medial side of the joint (Fig 1A-C, 3-4, arrowhead). Anaesthesia and muscle ablation also caused flattening and thickening of the anterior tip of the MC, close to the mandibular symphysis (Fig 1B3, C3, double headed arrow). Following removal from MS222, treated larvae moved their jaws significantly more frequently than control larvae (Fig 1D), potentially due to mild hypoxia. Although all AM-ablated larvae resumed movement, indicating regeneration, they had significantly fewer jaw movements than controls (Fig 1D) (S1 Fig), likely due to the reduced contractile ability of the regenerating fibres at the stages observed.

**Fig 1.**
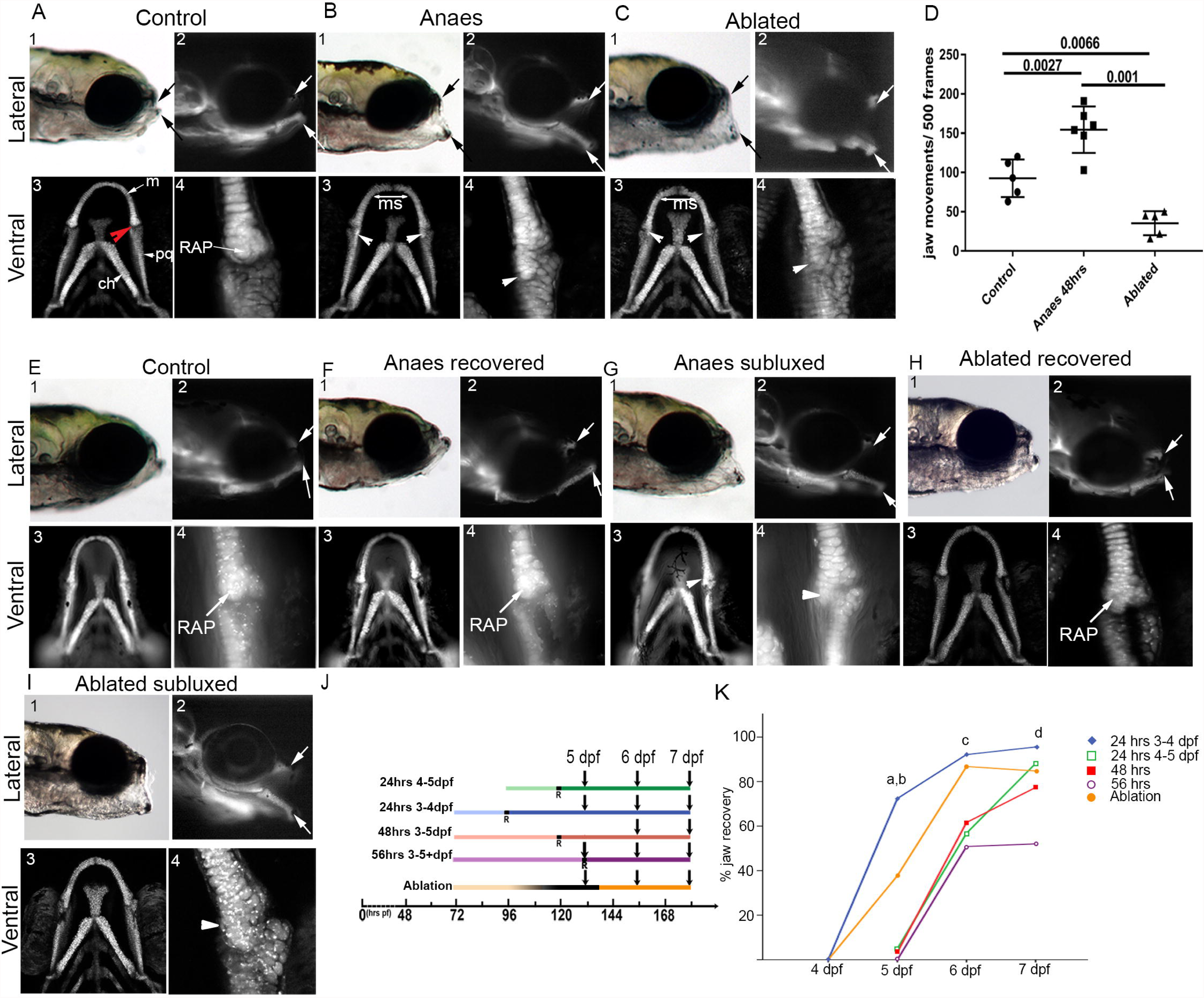
Restoration of muscle contraction is sufficient to drive the recovery of jaw function and shape. Zebrafish expressing Tg(*Col2a1aBAC:mcherry*) were imaged laterally (1,2) and ventrally (3-4) at 5dpf (A-C) and 7 dpf (E-I). Immobilisation and ablation caused joint subluxation (arrows, B1-2, C1-2) and flattening of mandibular symphysis (ms, B3-4, C3-4). The number of mouth openings per 500 frames at 5dpf in anaesthetised larvae, after wash out, and ablated larvae (D) analysed by 2-way anova. 7dpf anaesthetised and ablated larva subdivided by jaw position (F-I). Time course of percentage recovery following varying anaesthesia treatment analysed using Kruskal-Wallis test (K). (a) p=0.009 “24hrs 3-4” vs “48 hrs”, (b) p=0.008 “24 hrs 3-4” vs “24 hrs 4-5”, (c) p=0.049 “24hrs 3-4” vs “56 hrs” and (d) p=0.002 “24hrs 3-4” vs “56 hrs”. m; Meckel’s cartilage, ch; ceratohyal, pq; palatoquadrate, RAP; retro-articular process.

By 7dpf the majority of immobilised and of ablated larvae recovered from the subluxed position, such that they could now fully close the mouth (Fig 1E-I,1,2 arrows). Larvae with a “recovered” jaw position (Fig 1F, H) displayed complementary joint surfaces, and an appropriately positioned retro-articular process (RAP) (Fig 1F,H), resembling the appearance of unmanipulated controls. The remaining larvae retained their subluxed jaw position (Fig 1 G1-2, I1-2, arrows) with overlapping joint surfaces (Fig 1 G3-4, I3-4 arrowhead).

Severity of joint malformation correlates with timing and duration of anaesthesia treatment [5]. We varied the timing and duration of treatment to determine if these factors also determine ability to recover joint shape and function following resumption of movement. We immobilised zebrafish for 24, 48 or 56 hours from 72 or 96hpf (Fig 1J) and monitored the percentage of larvae that recovered following drug wash out and resumption of muscle activity (Fig 1K). Recovery was significantly affected by the timing of treatment. When movement resumed prior to 4dpf 95% of larvae recovered by 7dpf, a significantly higher percentage than the worst performing treatment group (those larvae immobilised for 56 hours) from which only 69 % recovered (Fig 1K). Folllowing ablation 84% of larvae recovered, however, due to low sample number (n=30) these were not included in statistical tests. These results indicate that there is sufficient joint plasticity at 5dpf to enable most larvae to recover functionality, however earlier resumption of movement significantly improves chance of recovery.

### Anaesthesia significantly alters the overall shape of the Meckel’s cartilage

To investigate the impact of immobilisation and restored movement on MC shape we used geometric morphometric analysis. In addition to immobilised and ablated larvae we included Myod-/- which lack jaw musculature [31] to model continued jaw joint paralysis. Control, Myod-/-, larvae anaesthetised from 3-5dpf and larvae with ablated AM were imaged live at 5dpf then each larva re-imaged at 7dpf. At 7dpf immobilised and AM-ablated larvae were classified as “Subluxed” or “Recovered”. Outlines of the MC at 5dpf (Fig 2A) were analysed using npManova and between-groups Principal Component Analysis (bgPCA) [16] to quantify how MC shape differs between the groups. Each principal component (PC) describes a trend in how the shapes within the analysis vary. Thus, similar shapes occupy similar positions along a PC axis while different shapes occupy opposing ends. A shape is described fully by all components as they may be similar in one component but differ in another.

**Fig 2.**
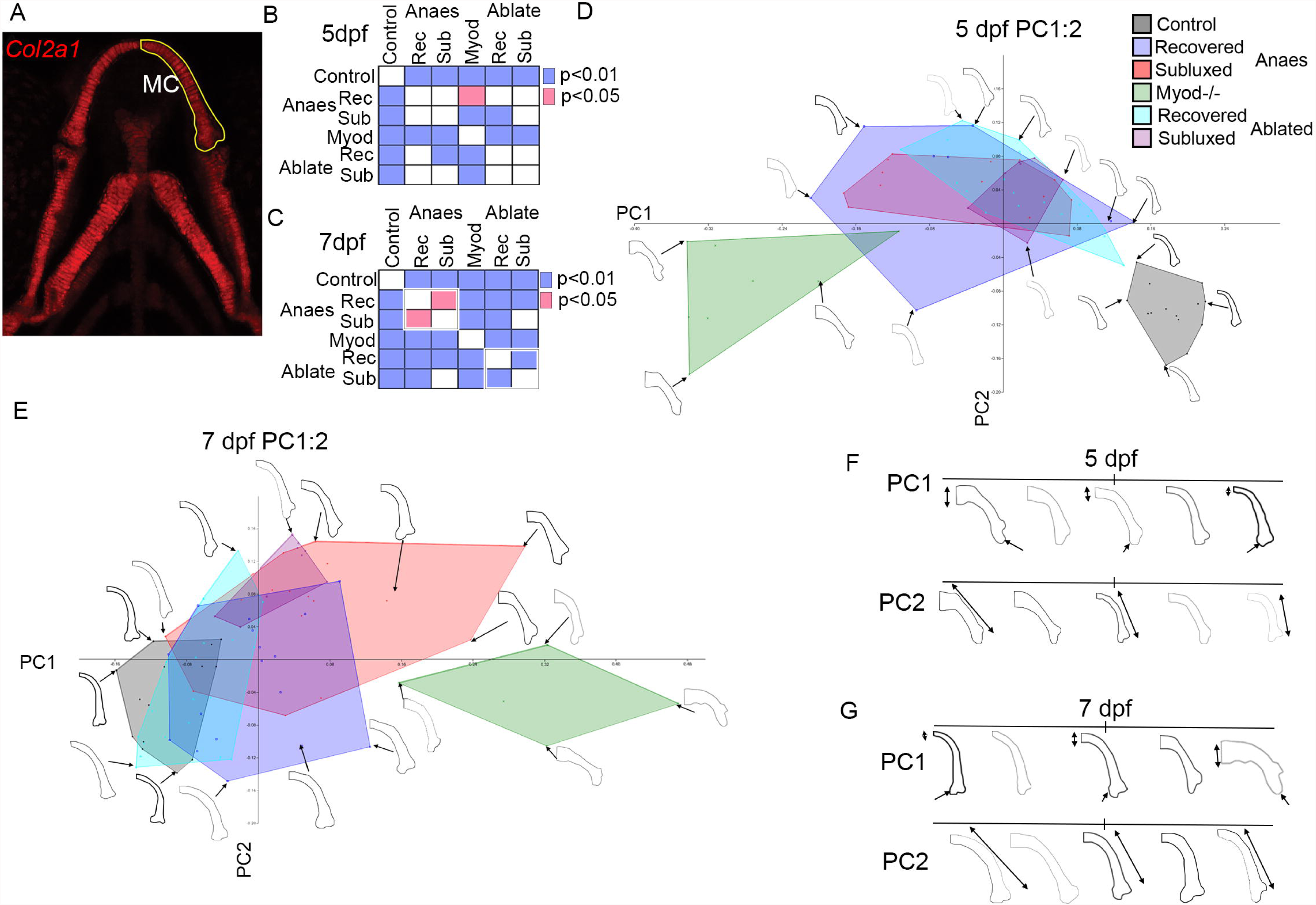
Shape cannot be used to predict recovery in anaesthetised or ablated larvae however restored movement significantly altered the shape of the Meckel’s cartilage by 7dpf. Graphic representation of results of npManova (B, C) of 2D outline of Tg(*Col2a1aBac:mcherry*) larvae (A). Principal component analysis of 5 (D) and 7dpf (E) treatment grouped by colour and subdivided by jaw position at 7dpf. Selected joint examples arranged according to PC1 and 2 at 5 dpf (F) and 7dpf (G) showing the increasing thickness of the MC (double arrow), overlapping joint shape (arrow) and altered curvature of the cartilage (PC2, double arrow).

Loss of muscle activity had a significant effect on the shape of the MC at 5dpf. The anaesthetised, ablated and Myod-/- larvae were all significantly different from control larvae (Fig 2B, S1 Table) and occupied different quadrants in PC1 and PC2 of the morphospace relative to control. PC1 accounts for 78.4% of variation in shape, separates the groups based on increasing MC width near the mandibular symphysis and degree of joint overlap (Fig 2D, F). PC2 accounts for 15.5% of the variation and captures the shape of the MC arch; control and Myod mutants are more gently arched than anaesthetised or ablated larvae. Ablated and anaesthetised larvae collocate while Myod mutants are dramatically different to controls. At 5dpf there was no significant difference in shape between larvae that will recover and those that remain subluxed at 7dpf (Fig 2B, D, F S1 Table). As such, MC shape at 5dpf does not reliably predict likelihood of recovery.

Resuming movement had a significant impact on the shape of the larval jaw at 7dpf. PC1 and PC2 accounted for 73.4% and 14.3% of variation between the groups; describing similar trends as at 5dpf (Fig 2E). The shapes of recovered jaws at 7dpf were significantly different from those that remained subluxed (Fig 2C, S2 Table) occupying an intermediate position in the morphospace plot, overlapping both the control and subluxed domains (Fig 2E, G). These data indicate that within a pool of similarly treated larvae a subgroup can recover a functional joint and the resultant recovered shape differs significantly from the shape of those that remain subluxed.

### Differences in cell orientation, shape and size precede recovery

Morphospace analysis highlighted the impact of altered load on mandibular symphysis and jaw joint shape. The differentiation of cells at both sites is affected by loss of muscle activity [5,15,16]. As chondrocytes mature their appearance changes from small spherical cells to larger ellipsoids [32]. Therefore, chondrocyte differentiation state can be inferred by measuring their size and circularity. A perfect circle has a circularity of 1, with decreasing values for elongated cells. We compared the shape and size of chondrocytes in the anterior MC in control, Myod-/-, anaesthetised and ablated larvae, with the latter two groups subdivided by their status at 7dpf. Cells flanking the mandibular symphysis were outlined and circularity and area calculated for each cell (Fig 3A-J). Due to the small number of AM ablated larvae that remained subluxed all further analysis on AM-ablated larvae were carried out on jaws that were classed as recovered at 7dpf. At 5dpf midline chondrocytes were significantly smaller and rounder in Myod-/- mutants and larvae that remain subluxed than controls and larvae that recover. At 7dpf only chondrocytes in larvae that remain immobile, i.e. Myod-/-, were significantly rounder than control, suggesting that movement drives maturation of this cell population.

**Fig 3.**
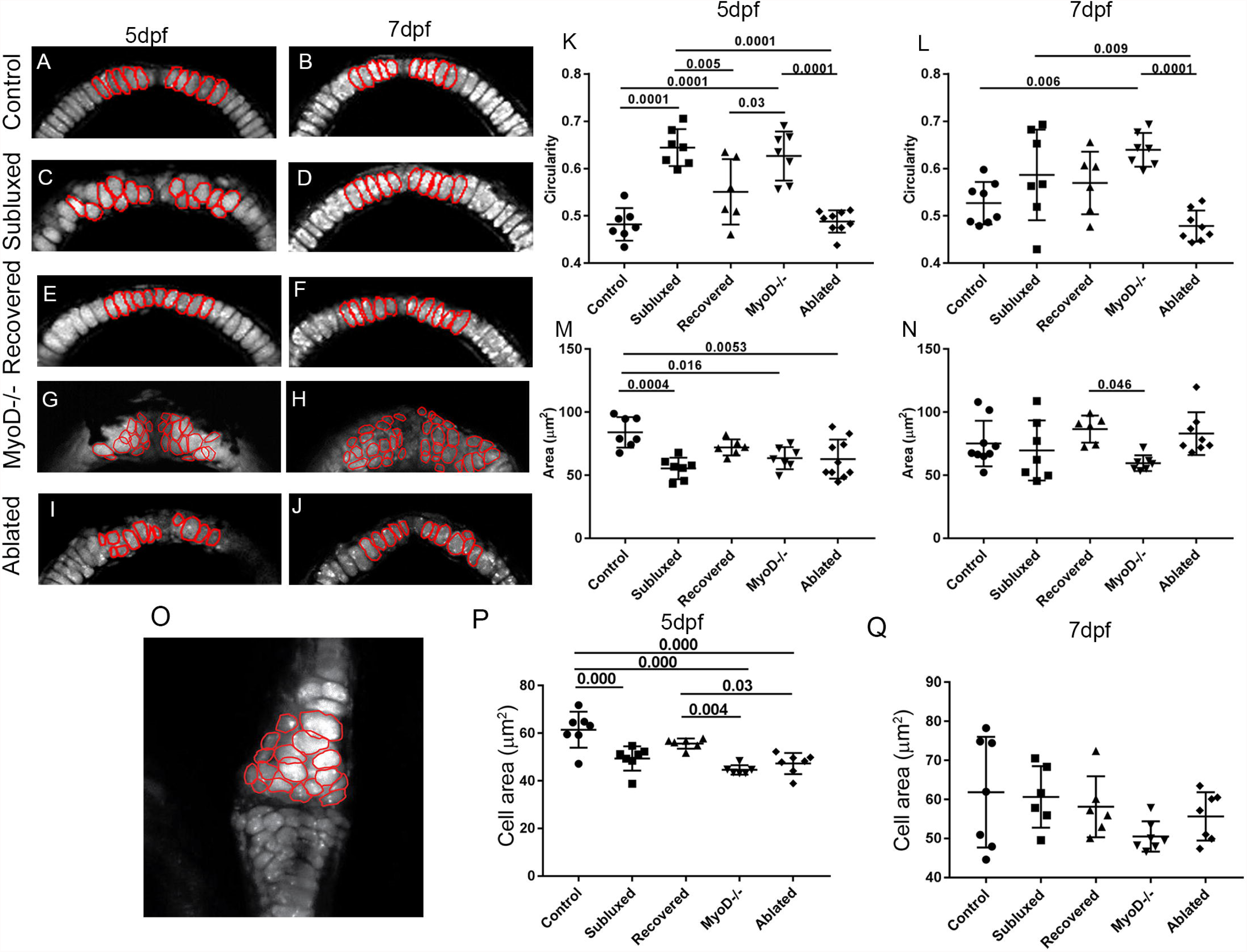
Chondrocyte maturation is less affected in the mandibular symphysis and jaw joint of larvae that recover. Confocal images of the mandibular symphysis of 5dpf and 7dpf control, myod-/-, anaesthetised and ablated zebrafish cartilage visualised using the Tg(Col2a1aBAC:mcherry) transgenic line (A-J). Confocal image of a representative jaw joint (O). Cell circularity (K, L) and cell area (M, N, P, Q) of 10 cells, outlined in red, between the insertion of the intermandibularis muscle (n=6 per condition) (A-N) and below intercalation in the joint (O). 1.00 represents a perfect circle.

Immobility leads to alterations to joint cell orientation and size [5]. To determine whether restoration of normal cell size and orientation is achieved during recovery we measured the chondrocytes located at the joint (Fig 3O-Q, S2 Fig). At 5dpf joint chondrocytes in anaesthetised larvae that will recover were statistically indistinguishable to controls; while chondrocytes in those that failed to recover, Myod-/- and AM-ablated larvae, were significantly smaller than the chondorcytes of controls (Fig 3P). No significant difference in cell size was seen between any of the groups at 7dpf; due to the variability of control cell size (Fig 3Q). Analysis of the orientation of the major axis of joint cells relative to the palatoquadrate surface, revealed significant differences in orientation of cells at 5dpf on both the medial and lateral joint sides in larvae that fail to recover relative to controls, whereas joint cells from larvae that will recover do not differ from controls. By 7dpf the orientation of cells in subluxed treated larvae are no longer significantly different from controls and recovered larvae, whereas Myod-/- larvae, which receive no mechanical input remain significantly different to controls (S2 Fig B, C). This data demonstrates that variation in size and orientation of cells located at the joint and symphysis at 5dpf are associated with ability to recover; and that reapplication of mechanical stimulus allows cells to reorientate.

### Movement drives restored expression of joint associated genes

Loss of muscle activity in amniotes leads to cartilage fusion in the joint territory driven by loss of joint specific markers in favour of the expression of *Collagen 2* (*Col2*) and other cartilage markers [6,12,33]. We used the relationship between *Col2a1*, a marker of mature chondrocytes, and *Sox10*, a marker of neural crest [34] expressed in immature craniofacial chondrocytes and joint precursor cells (Fig 4A), to determine if joint boundaries re-emerge during recovery. At 3dpf cells expressing *Sox10* were seen across the whole joint, with particularly strong expression on the medial side where the RAP forms (Fig 4A 2, 3, 6). At 5 and 7dpf cells in the joint line, tip of the RAP and 2-4 cells flanking the mandibular symphysis express high levels of *Sox10* and much lower levels of *Col2a1* (Fig 4 A4, 5, 7).

**Fig 4.**
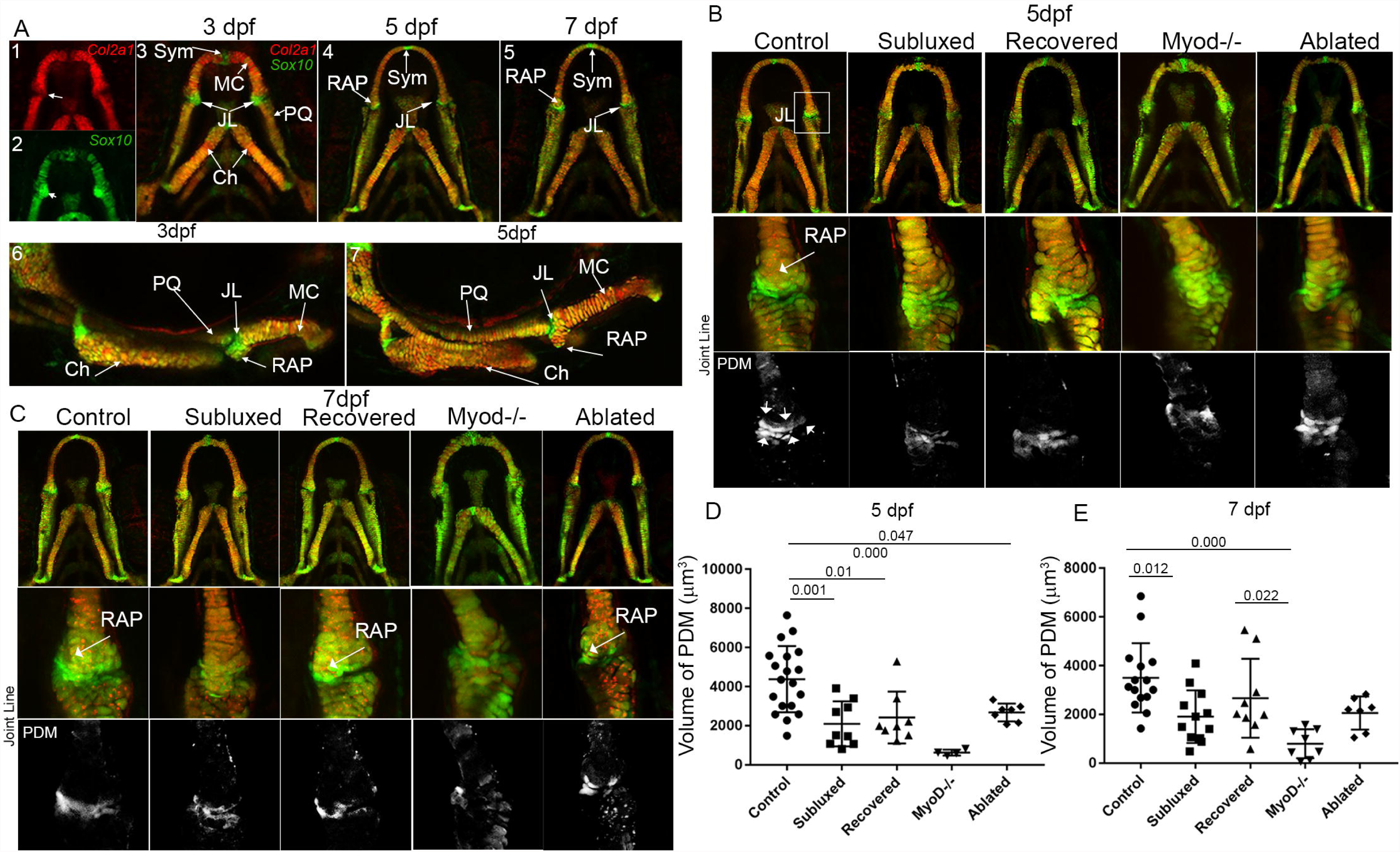
Restored movement drives the restoration of the joint line. Ventrally imaged zebrafish expressing Tg*(Col2a1aBAC:mcherry)* and *Tg(-4725sox10:GFP)*^*ba*4^ at 3, 5 and 7dpf and laterally at 3 and 5dpf. High magnification confocal images of the joint line of 5dpf (B) and 7dpf (C). Ablated and anaesthetised larvae which were subdivided by their jaw position at 7dpf. Joint volume (D, E) derived from the “product of the difference of the mean” (PDM) of the joint (white box). JL; joint line, Syml; mandibular symphysis PQ; palatoquadrate, mc; Meckel’s cartilage, Ch; ceratohyal.

To alleviate variation in the intensity of *Sox10* or *Col2a1* due to copy number variation or imaging parameters we extracted the anti-correlation component of the “product of the difference of the mean” (PDM) [35] of the confocal dataset. In 5dpf controls *Sox10* was expressed across the joint line in cells where *Col2a1* was not expressed; these cells were clearly identifiable in the PDM (Fig 4B, arrows). In comparison immobilisation lead to a significant reduction in the size and definition of the joint domain, particularly apparent in Myod mutants (Fig 4B, D). The magnitude of the anti-correlation component, indicative of how different the two channels were, was also significantly reduced in Myod-/- larvae at 5dpf (S3 Fig).

At 7dpf the relative expression of *Sox10* and *Col2a1* remained significantly disrupted in subluxed and Myod-/- larvae (Fig 4C, E), whereas the joint region of recovered larvae and AM-ablated larvae were no longer significantly different to controls (Fig 4E). Thus, larvae capable of recovery reacquire a group of cells with high *Sox10* low *Col2a1*, suggestive of immaturity/plasticity in the joint, whereas larvae that fail to recover or remain paralysed fail to maintain these joint specific cells.

### Recovery is driven by changes to cell proliferation and migration

We hypothesised that proliferation and/or migration of these *Sox10* high *Col2* low cells might drive joint recovery. Cell proliferation following the resumption of movement was measured by treating larvae with BrdU from 5-7dpf, at which time the larvae were sorted into recovered and subluxed groups, stained immune-fluorescently and quantified (Fig 5C, D). BrdU incorporation was significantly increased, relative to control, on both the medial and lateral side of the joint in subluxed larvae. In contrast, proliferation was only increased on the lateral side of the joint in recovered larvae (Fig 5 D). BrdU+ cells were present in larger numbers in the lateral joint margin (Fig 5A arrowhead) in both groups relative to control.

**Fig 5.**
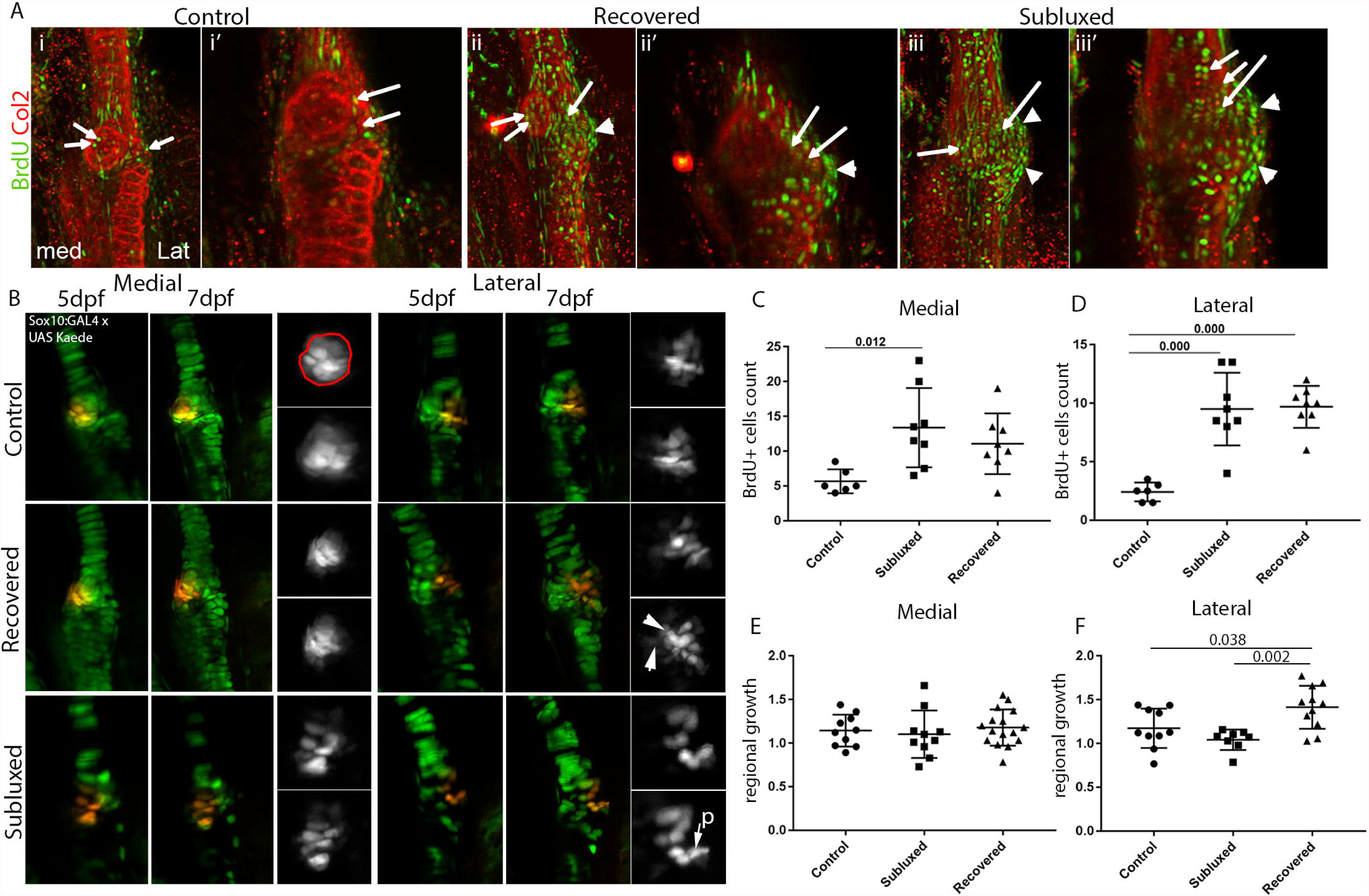
Cell proliferation and migration is differentially regulated in zebrafish that recover relative to those that remain subluxed. BrdU incorporation (green) between 5-7dpf (A). Arrows show BrdU+ cells within the cartilage. Arrowhead shows BrdU+ incorporation outside the cartilage. Cells expressing Tg(Sox10:GAL4-VP16) and Tg(UAS:Kaede) are green until photoconverted. Red photoconverted cells located on the medial or lateral sides of the joint at 5dpf (left panels), reimaged at 7dpf (middle) with detail of photoconverted cells (right). Arrowhead indicates migration, p indicates proliferation. Number BrdU+ cells on the medial and lateral side of the joint in control, anaesthetised recovered and subluxed zebrafish. Fold change in total area of cells (red outline in B) expressing photoconverted red kaede in control, anaesthetised recovered and subluxed. The impact of side and treatment on BrdU+ counts and Kaede growth analysed by MANOVA.

Cell migration during recovery was assayed by photoconvertion of kaede-expressing cells in zebrafish carrying Tg(Sox10:GAL4-VP16) and Tg(UAS:Kaede) transgenes. Kaede protein is green in its native state but is irreversibly photoconverted to red using UV light. 8-10 cells were photoconverted at 5dpf on either the medial or the lateral side of joints in larvae that were anaesthetised and in controls. These larvae were then reimaged at 7dpf to track the behaviour of the photoconverted cells. The spread of cells across the joint was calculated as the change in the area of the photoconverted domain (Fig 5E, F), measured from max projections between 5 and 7dpf (Fig 5B, red outline). Cellular migration was significantly increased on the lateral side of recovered but not subluxed larvae (Fig 5E, F). There was also evidence of cellular reorientation on the lateral side of recovered joints that was not apparent in subluxed larvae (Fig 5B arrowhead). In comparison, no changes to cellular orientation or spread of photoconverted cells were detected on the medial side. This data indicates that cells at the joint retain the capacity to adapt to the mechanical environment and sculpt the joint by modifying regional patterns of cell migration and proliferation.

### Joint associated structures such as ligaments and their insertions are restored in recovered larvae

Immobilisation during development has been shown to have a significant impact on the structure of joint associated tissues such as ligaments and their attachments [6,11,36]. The RAP, the insertion site of the interoperculomandibular ligament (IOM) is significantly impacted by loss of mechanical load (Fig 1,2) [5,16]. C*ol10a1:citrine (ColX)* was expressed by osteoblasts in the ligament entheses (Fig 6A,E) [37] and at a lower level throughout the IOM ligament (Fig 6A, E). It was used to assess the impact of immobilisation (Fig 6 A-D) and recovery (Fig 6E-I) on osteoblast number in the RAP and ligament width. Immobilisation caused a significant reduction in osteoblast number in the RAP at 5dpf (Fig 6C). By 7dpf the number of osteoblasts at the RAP of subluxed, but not recovered larvae, were significantly fewer than controls (Fig 6 E-H).

**Fig 6.**
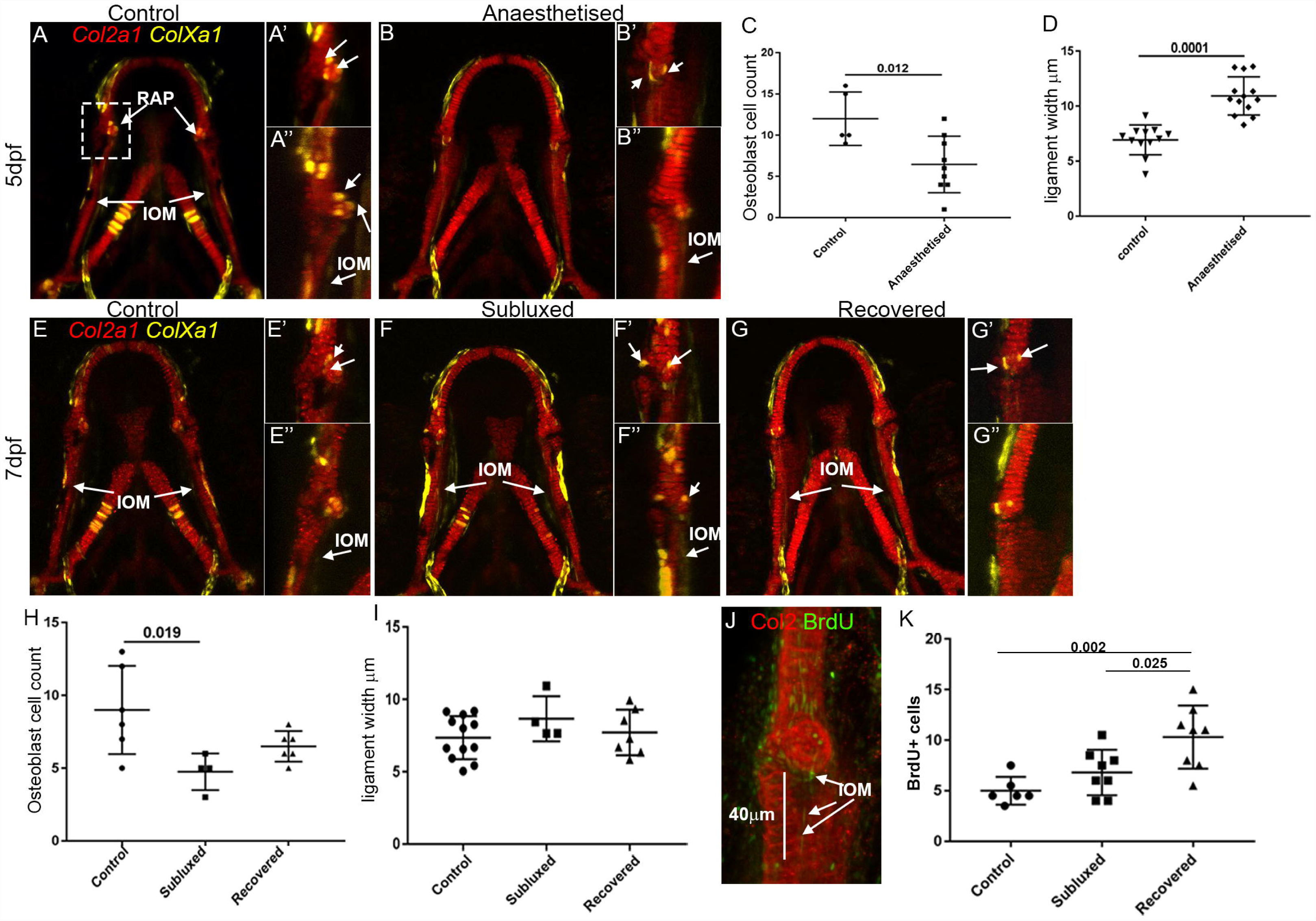
Recovery restores joint associated structures. Confocal images of control and anaesthetised TgBAC(Col10a1a:Citrine)^hu7050^ and Tg*(Col2a1aBAC:mcherry)* were imaged ventrally at 5dfp (A-B”) and again at 7dpf (A-G”). At 7dpf they were split into recovered and subluxed by jaw position. Detail images of RAP (‘) and RAP rotated approximately 90° (”)Counts of ColX expressing osteoblasts located at the RAP (arrows in detail images) at 5dpf (C) and 7dpf (H). Width of the Interoperculomandibular ligament (IOM), measured at 3 sites along the ligament at 5dpf (D) and 7dpf (I). Counts of BrdU incorporation into the IOM between 5dpf and 7dpf, prepared within 40μm of its insertion into the RAP in control and anaesthetised larvae split by jaw position at 7dpf (J-K). Analysed by Anova.

At 5dpf, the width of the IOM ligament was significantly increased in immobilised larvae reflecting a loss of the tight cohesive structure characteristic of ligaments (Fig 5D). At 7dpf the subluxed cohort remained marginally larger but this was not significant, possibly due to the small sample number (n=4). We verified the recovery of the ligament by comparing the number of ligament cells that incorporated BrdU between 5 and 7dpf in control, subluxed and recovered larvae. Larvae that recover proliferated significantly more between 5 and 7dpf than either control or subluxed larvae. Therefore, larvae that successfully recover a functional shape also repair the degenerative impact of immobilisation on the interzone and joint associated tissues such as ligaments.

### Recovery potential is associated with regional differences in the pattern, magnitude and orientation of mechanical stimuli within the jaw

Mechanical forces act as a form of positional information with cells behaving in a regionally specific manner dependent on its mechanical environment. We hypothesised that joint shape, cellular orientation and differentiation could feed into the mechanical environment, which in turn regulates the cells, producing a feedback loop resulting either in success or in failure. Renewed movement is required for the initiation of shape recovery and may drive the rapid correction seen in recovered larvae. Finite element (FE) models of representative 5dpf control, anaesthetised recovered and subluxed larva were analysed to identify potential differences in the pattern of biophysical stimuli experienced by larvae that recover and those that remain subluxed. The range of motion of the tip of the jaw in control and anaesthetised larvae was 53.3 (+/- 18.4) μm and 57.71(+/- 22.5) μm respectively, which was replicated by the displacement of the FE models during the closure step (Fig 7A-C). The insertion of the IOM (interosseous membrane) ligament at the jaw joint generates a distinct peak in strain, which is located at the tip of the RAP in controls (Fig 7D, J, Fig S6A). This peak correlates with the position of *Col10a1* expressing osteoblasts; which are mechanically responsive and reduced in anaesthetised larvae (Fig 6). The mandibular symphysis is also a site of increased mechanical load in models of 5dpf control and recovered larvae (Fig7, D-E,J-K arrow head) but not the subluxed model. The orientation/direction of the principal strains within a tissue are important regulators of cell orientation. In the MC the orientation of maximum principal strain, as indicated by the direction of the arrows (Fig 7G-I), closely follows the curvature of the cartilage with a distinct peak at symphysis. Thus cells within the MC are orientated perpendicular to these strains. While the magnitude of the loads are reduced, relative to control, it is still possible to identify a distinct peak at the symphysis in the model of recovery which is absent in the subluxed (Fig 7I).

**Fig 7.**
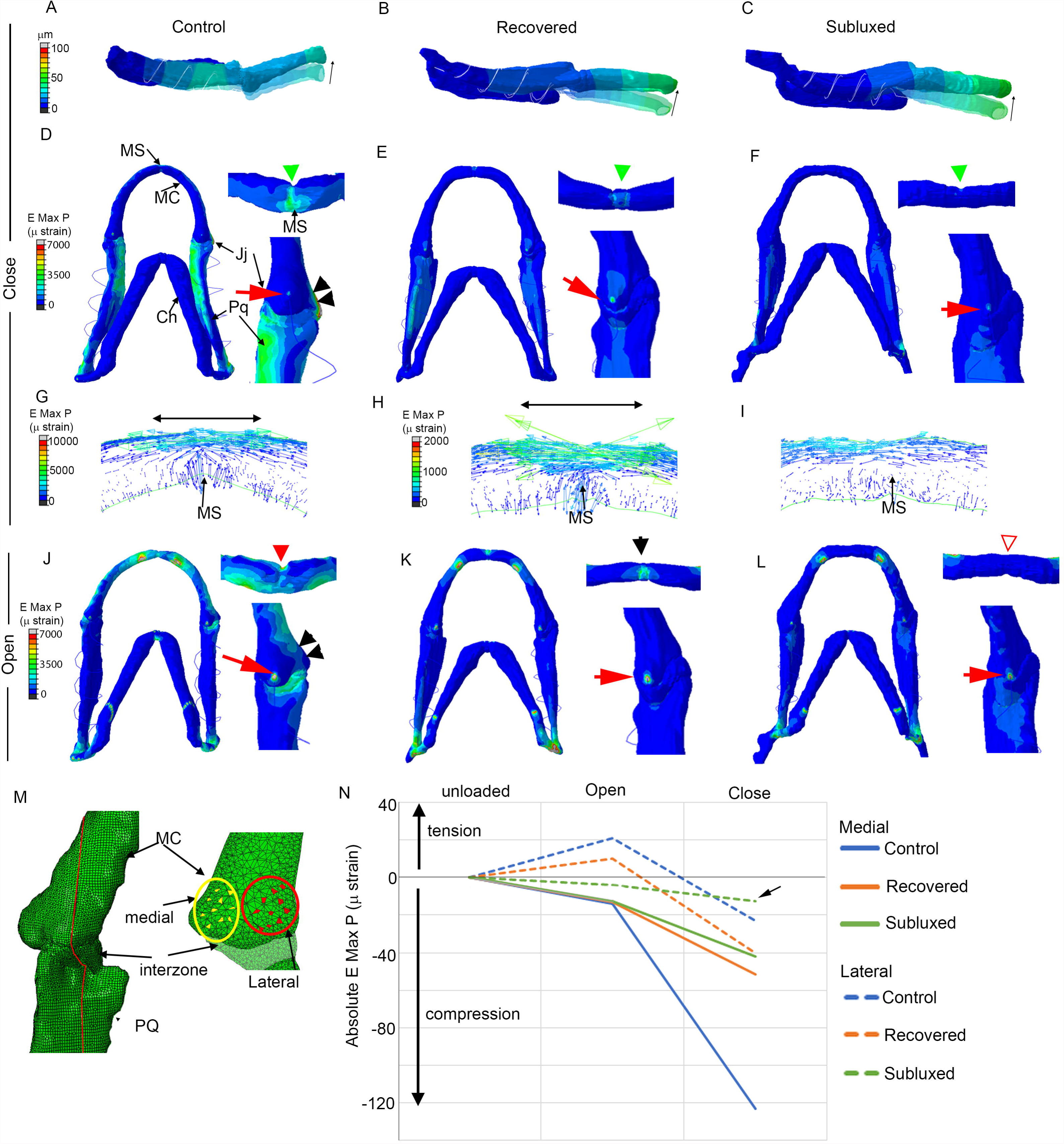
Finite element (FE) analysis reveals distinct differences between the patterns of biophysical stimuli. FE model simulating jaw displacement during closure in models of a 5 dpf control and anaesthetised larvae that recover (B) or remain subluxed (C). Displacement is shown as the block colour model relative to the semi-transparaent model of the jaw original position, with the direction of motion indicated as an arrow. Finite element models of maximum principal strain created from confocal stacks of 5 dpf control (D,G, J) and anaesthetised larvae that recovered (E,H, K) or remained subluxed (F,I,L) during closure (D-I) and opening (J-L). Each panel shows a ventral view, enlarged image of the joint (Jj) and mandibular symphysis (MS). Strain was elevated at the symphysis (green arrowhead), RAP (red arrow) and on the lateral side of the joint (black arrow heads) (D-F, J-L). The orientation and magnitude of the strains are indicated by the direction, size and colour of arrows in the plane through the mandibular symphysis (G-I). Graph of the average absolute maximum principal strain during opening and closure (N), extracted from 10 elements on the medial and lateral side of the joint in a plane located at the midpoint of the joint (redline M).

We identified distinct differences in the regional behaviour of cells between larvae that remain subluxed and those that successfully recover. FE modelling has been shown to predict changes to joint shape and cellular behaviour in the developing jaw joint [5]. We used the FE models to identify if larvae that recover also exhibit distinctly different patterns of biophysical stimuli. In controls, the medial side of the joint is highly loaded (Fig 7D, J, black arrowhead). Recovery is generated by a specific pattern of cell proliferation and migration that varies medial-laterally. In joints that recover there is a significant increase in both proliferation and migration on the lateral side of recovering joints that is absent from those that remain subluxed. To determine if the magnitudes and pattern of loading differed between regions of the model we extracted the absolute maximum principal strain for 10 nodes on the medial and lateral side of the joint (Fig 7M) and graphed the results (Fig 7N). In all three models the medial side experiences variable but negative absolute maximum strain indicating a dynamic compressive environment. In comparison, the lateral side of both control and recovered models oscillates between positive (tensional) and negative (compressive) strain while the subluxed surface remains under a lower magnitude negative (compressive) strain.

## Discussion

Here, we present a novel zebrafish model of developmental joint dysplasia; characterising the cellular mechanisms underpinning the recovery of joint function in malformed joints. Immobilised joints lack the complex interlocking joint shape characteristic of synovial joints, which maintain joint position and range of motion, therefore recapitulating many of the features of DDH [8]. Here we demonstrate that the resumption of movement in immobilised zebrafish joints can be used to examine the impact of suboptimal joint shape on joint health, raising the prospect of following manipulated joints throughout the life of the individual.

The zebrafish jaw joint, between the MC and Palatoquadrate, was the focus of this study. This joint is synovial and can develop osteoarthritis [38] and has all the key features of mammalian skeleton including articular cartilage, joint cavity, synovium, tendon and ligaments [39,40]. The zebrafish also has a growing array of genetic tools [41] and is amenable to live imaging making it ideal for longitudinal studies. Tricaine immobilisation alters the shape of the jaw joint and MC by altering cell differentiation and proliferation [5,15] and is similar to the method used in the chick model of DDH [10]. Laser muscle ablation induced a similarly malformed jaw joint without the systemic issues associated with drug treatments. We found a significant proportion of both the anaesthetised and ablated larvae could recover a normal jaw position and approximately normal joint shape while those that fail to recover remain subluxed and have a significantly different jaw shape, reflecting the variability seen in human cases of joint dysplasia.

DDH symptoms can range from a mild joint laxity that can resolve itself, up to full subluxation requiring reduction. Prognosis is correlated with severity of the malformation of the acetabulum and age of treatment [1]. We hypothesised, that within a similarly treated group, jaw joint shape would be an indicator of recovery. However, morphometric analysis indicated that the shape of the MC at 5dpf was a relatively poor indicator of likelihood of recovery. As a simplified 2D analysis was performed, it is possible that 3D aspects of the shape that might influence recovery were not captured by the analysis. The relationship between the shape and position of the palatoquadrate surface relative to the MC could play a substantial role in recovery and wasn’t included in the morphometric analysis.

While shape alone could not predict which larvae would recover, chondrocyte behaviours at the mandibular symphysis and joint could. The orientation of cells in the joint is significantly disrupted by immobilisation [5]. At 5dpf, cells in the mandibular symphysis and jaw joint of larvae that remain subluxed were rounder and less differentiated than those that recover. Interestingly, by 7dpf cell orientation is corrected in all samples bar the Myod mutants. The larval response to immobilisation and recovery is a spectrum determined by differences in cell behaviour.

Embryonic immobilisation is associated with loss of the articular surfaces, joint fusion and the degeneration of structures such as meniscus and patella [33]. 5dpf immobilised larvae lost the definition of the interzone, which gives rise to the articular cartilage [42]. By 7dpf the anti-correlative relationship between *Sox10* and *Col2a1* expression in the interzone of recovered larva was redefined but remained abnormal in joints that fail to recover (subluxed and Myod mutants). The development of ligaments [6,33] and their insertion sites (Entheses) are particularly vulnerable to loss of mechanical stimulus [43]. In immobilised and ablated larvae at 5dpf the interoperculomandibular ligament was wider and looser and ossification at the entheses was significantly reduced. By 7dpf recovery ameliorated this impact significantly. Thus, movement corrects not only the shape of the joint but also drives the recovery of vital joint associated structures.

Skeletal morphogenesis requires preferential regional growth, a feature lost during immobilisation. It is achieved by the local moderation of cell migration, proliferation, cell shape and matrix synthesis [33]. Cellular reorientation and shape change are a feature of recovery however immobilisation has been shown to alter patterns of cell proliferation [7,44]. BrdU incorporation and Kaede photoconversion identified a specific pattern of cell proliferation, migration and growth on the lateral side of the joint that was only present in larvae that recover. These local differences in cell behaviour may be driven by specific biomechanical differences between recovered and subluxed larvae.

We developed Finite Element (FE) models based on 3D confocal datasets of control and anaesthetised larvae at 5dpf. The two chosen anaesthetised larvae were representative examples of those that recover and those that remain subluxed. We identified variations between the mechanical stimuli experienced by these anaesthetised larvae, which may in turn drive the restoration of normal joint shape and function. FE results suggest that cells in the Meckel’s cartilage orientate themselves perpendicular to the main direction of strain and that this signal is stronger in larvae that successfully recover. This is in keeping with reports that cells align nearly perpendicular to dynamically varying stresses [45]. The mechanoregulation of chondro-progenitor and chondryocyte proliferation and biosynthesis has been extensively studied in a variety of culture system including explants and monolayer in response to a wide range of mechanical stimuli including compression, tension, hydrostatic pressure and fluid flow [46]. The response of cells to a particular load is dependent on the type of load, its mode of application i.e static or cyclic and the magnitude of this load. Cyclical loading of chondrocytes cultured in alginate beads had a positive effect on chondrocyte metabolism increasing cartilage matrix synthesis while static load was degenerative[47]. Cyclical loading impacts the differentiation of cells, increasing Indian hedgehog expression and stimulating chondrocyte proliferation[19]. Stretch also stimulated the proliferation of immature chondrocytes, but not that of hypertrophic chondrocytes [48]. Ultimately load appears to differentially regulate the behaviour of cells[18]. Thus morphogenesis is regulated through a combination of intrinsic regulatory signals produced by the tissue itself e.g. Wnt or Bmp signalling, and extrinsic signals produced by mechanical stimuli. These extrinsic stimuli will in turn alter the expression of intrinsic regulators creating a vicious circle of degeneration amplifying the original malformations. Thus the difference between success and failure may be initially quite minor but could be exacerbated by this “intrinsic-extrinsic” feedback loop. It also demonstrates that successful recovery is very dependent on the individual, their specific cellular orientation, joint shape and mechanical environment. This is corroborated by the long term prognosis of DDH which even after treatment is variable[49].

It is possible that higher vertebrates, which lack the regenerative capacity of zebrafish [50], would not respond as dramatically to the resumption of movement, though even in humans cases of DDH can resolve spontaneously with no intervention [1]. As skeletal plasticity is preserved in higher vertebrates we believe the zebrafish model could provide useful insight into the mechanism regulating this plasticity. Using either anaesthesia or muscle ablation it will be possible to target specific windows of plasticity during joint development and identify what processes are disrupted and ultimately determine how some larvae are better positioned for recovery, in the hope of identifying why the prognosis from some DDH patients is so severe.

## Materials and Methods

### Zebrafish husbandry and lines

Zebrafish were housed as previously described [51]. Animal experiments were approved by Bristol University animal ethics committee and the UK Home Office. All transgenic and mutant lines have been previously described: TgBAC(*col2a1a:mCherry*)^hu59^ [52]; Myod^fh261^ [31]; *Tg(-4725sox10:GFP)^ba^*^4^ [53]; *symhc:EGFP* [54]; Tg(*Sox10:Gal4-VP16*) [55]; Tg(UAS:Kaede) [56]; TgBAC(col10a1a:Citrine)^hu7050^ [57]. Sample numbers used are listed in supplementary S 3 Table.

### Pharmacological treatment and imaging

Fish were treated for 24 hours beginning 72 hours post fertilisation (hpf) and 96hpf, 48 hrs from 72hpf, or 56 hours from 72hpf (Fig 1E) with 0.1mg/ml MS222 (Tricaine methanesulfonate)(Sigma), diluted in Danieau’s buffer refreshed twice daily. Jaw position was assessed daily between 5 and 7dpf (Fig. 1C) and the percentage recovery calculated from at least 4 plates of 30 larvae. Tracked cohorts were imaged on 5 and 7dpf.

### Live imaging

Larvae were sedated, embedded ventrally in 0.5% LMP agarose (Sigma) and fluorescence visualised on a Leica SP8 system using a 10x objective. After imaging larvae recovered in fresh Danieau’s buffer.

### Muscle Ablation

The Tg(*smyhc:EGFP*) transgenic line was used to visualise muscle position. Anaesthetised 3dpf larvae were mounted ventrally in 0.2% LMP agarose in Danieau’s and adductor mandibulae (AM) muscles bilaterally ablated with a 440nm nitrogen Micropoint laser connected to a Zeiss Axioplan microscope. Ablation success was confirmed by imaging after ablation. As *smyhc:eGFP* is expressed in slow fibres, the efficacy of laser ablation was verified in 3dpf fixed larvae using fluorescent immunohistochemistry [58] against GFP (ab13970 abcam, 1:200) and skeletal myosin (A4.1025 DSHB, 1:200) (S4 Fig)

### Jaw movement movies

Movies were taken as previously described [5]. Larvae were imaged at 30msec per frame for 500 frames and the number of mouth movements and jaw displacement was recorded. Results were analysed by Anova (SPSS).

### Analysis of shape variation

Changes to the Meckel’s cartilage (MC) shape were quantified using two-dimensional (2D) geometric morphometrics [16]. Outlines of the MC of 5dpf and 7dpf controls, anaesthetised and ablated larva were digitised and analysed by non-parametric MANOVA (npManova) and between-groups Principal Components Analysis (bgPCA). The anaesthetised and ablated larvae were subdivided by the functional position of their jaw (subluxed or recovered) at 7dpf.

### Cell orientation, area and shape

Separate Z-projections of 2-3 consecutive slices were created from *Col2a1* labelled images of the MC through the mandibular symphysis and jaw joint. Cell area and the circularity of a minimum of 10 cells between the insertion points of the intermandibularis anterior muscle were analysed [16]. Cell orientation and area were measured for a minimum of 6 joints per condition at 5 and 7dpf [5]. Chondrocytes were subdivided into those located within 3 cell widths of the medial or lateral side of the joint. Graphs were produced and analysed using PAST [59] and PRISM (Version 7).

### *Sox10 Col2a1* colocalisation

Larvae carrying *Col2a1:mcherry* and *sox10:GFP* transgenes were anaesthetised for 48hrs from 3dpf, allowed to recover, imaged at 5 and 7dpf and processed using a custom MATLAB tool. Stacks were binarised using a threshold to remove background noise. The product of difference from mean (PDM) of each image was calculated [35]. In this data positive values correspond to correlation and negative to anti-correlation between the expression of *Sox10* and *Col2a1*, after each has been normalised by average brightness. The anti-correlation dataset was transformed to a positive image, the joint region selected in 2D and the boundary projected across all slices of the image stack. Within each boundary, pixels above the intensity threshold were fit with an alpha shape [60] yielding a measure of the enclosed volume.

### Kaede protein photoconversion

Control and anaesthetised double transgenic Tg(*Sox10:GAL4-VP16*) Tg(UAS:Kaede) larvae were mounted ventrally at 5dpf and 8-10 Kaede expressing cells on the medial or lateral side of the joint were photoconverted [58]. Following photoconversion, larvae were kept individually, reimaged at 7dpf and jaw position assessed. Maximum projections of the red channel were prepared [61] and the area of the photoconverted region was calculated before and after recovery.

### BrdU immunostaining

Control and 48hrs anaesthetised larvae were treated with 3mM BrdU (Sigma) diluted in Danieau’s solution from 5-7dpf. BrdU incorporation was visualised by immunohistochemistry [58] using mouse anti-BrdU (Sigma, 1:100) and rabbit anti-collagen II (Abcam, 1:200), prior to imaging on confocal SP8. BrdU positive cells within the jaw joint cartilage, were counted and divided into the medial and lateral side of the joint.

### Finite Element Modelling

Representative confocal images of 5dpf recovered and subluxed larvae expressing *Col2a1:mcherry* were converted into 3D finite element models for comparison with previously published 5dpf control models using Hypermesh (Version 10, Altair Engineering) [5]. All models had a Young’s modulus of 1.1 MPa for the cartilage and 0.75MPa for the interzone and mandibular symphysis, and poisons ratio of 0.25 [as described in 5]. Muscle attachments were added to the FE-models for mouth closure (adductor mandibulae) and opening (protractor hyoideus and intermandibularis) using confocal datasets for reference for attachment sites (Fig S5 A,C,E, O-Q, Table S5). Spring elements were used to simulate the Interoperculomandibular ligament (Fig S5, O-Q, S4 Table) [7]. Anaesthesia impacts the function of both the muscle and ligaments. In the chick Immobilisation causes to a reduction in muscle size and its ability to transmit forces [62,63]. The magnitudes of the forces applied to the model were therefore adjusted to 75% of the normal estimation [derived from 63]. Immobilisation of the mouse cruciate ligament also reduced its stiffness and peak force[64]. The stiffness of the spring was reduced to 75% (1.3E-0.4 kN/mm). Immobilisation disrupts the definition of the interzone. While a clear interzone could be identified in control and recovered images it was not possible to identify a clear interzone spanning the joint space in the subluxed dataset, therefore the joint was modelled as a fusion with a collar of interzone (Fig S5 O). The geometrically linear models of approximately 2 million tetrahedral elements were imported from into Abaqus FE-software (v6.14 Simulia, Dassault Systèmes), the spring elements were assigned and the model was analysed (v6.10.2 Simulia, Dassault Systèmes).

## Acknowledgments

The authors would like to thank the Wolfson Bioimaging Centre, Jen Bright and David Gurevich for their technical advice and support.

## Supplementary Figures

S1 Fig. Muscle repair post ablation. *Smyhc:egfp* and *Col2a1:mcherry* transgenic zebrafish were bilaterally ablated at 3dpf and imaged at 4dpf and 5dpf. Yellow arrowhead indicate ablation site and repairing muscle.

S2 Fig. Cell orientations on the medial (A, green cells) and lateral (A, red cells) in wild type, anaesthetised recovered and subluxed and myod-/- and bilaterally ablated zebrafish jaw joints. Orientation angle, calculated from the longest axis of the cell (A, insert) of chondrocytes 5dpf (B) and 7dpf (C) zebrafish in the Meckel’s cartilage element of the joint, plotted on circular histograms (rose plots), where 0° lies on the medial side of the joint and 180° at the lateral side of the joint. n=6 joints per experimental condition (1 or 2 refers to number of joints per blue wedge). Histogram bins equal 20°. The red line marks mean orientation and the green line marks the 95% confidence interval and analysed by MANOVA.

S3 Fig. Differential expression of *Sox10:gfp* and *Col2a1:mcherry* is lost in the joints of Myod/- zebrafish. The intensity of product of the difference of the mean (PDM) between *Sox10:gfp* and *Col2a1:mcherry* double transgenic in control, anaesthetise recovered and subluxed, Myod mutants and double ablated zebra fish.

S4 Fig. Muscle fibres numbers reduced after ablation. Confocal images of control (A,C,E) and adductor mandibularis (AM) ablated (B,D,F) transgenic *smyhc;EGFP* (A-B) zebrafish following immunohistochemistry for total Myosin (C-D) show significant reduction in the reduction in the number of fibres (G). Percentage of Fish divided by the success of ablation (H). Arrowhead indicates adductor mandibularis.

S5 Fig. Building Finite element (FE) models.

3D surfaces were generated from confocal images of 5dpf *Smyhc:egfp* and *Col2a1:mcherry* control (A) and anaesthetised larvae (C,E) were used to generate FE models (O-Q). Anaesthetised larvae were subdivide by their jaw position at 7dpf (C, E). By 7dpf the recovered larva (F) was significantly different in shape to the subluxed (D). Sections through the jaw joint (J,K,N indicated by white line) show a continuous interzone in the joint (J’, N’, arrowhead) and mandibular symphysis (MS) (G, I, arrowhead) of controls and the recovered larva at 5dpf but not the subluxed. A spring modelled the Interoperculomandibular ligament (IOM) between the retro articular process (RAP) and Hyod bone (P-R, yellow line). 50% of maximum muscle loads (measured in Newtons (N)) were applied at the origin and insertion of the muscle The adductor mandibulae (AM) (O’,P’,Q’, blues line) was applied to model mouth closure. To capture the wider insertion of the intermandibularis (IM) and protractor hyoideus (PH) (O) the forces were applied across 10 nodes to model of mouth opening (O”-Q”). White dots indicate the area of applied constraints in either the x,y, or z orientation (O).

S6 Fig. Finite element (FE) models of minimum principal strain (Emin).

Finite element models of minimum principal strain (compressive) created from confocal stacks of 5 dpf control (A,D) and anaesthetised larvae that recovered (B, E) or remained subluxed (C,F) during closure (A-C) and opening (D-F). Each panel shows a ventral view, enlarged image of the joint (Jj) and mandibular symphysis (MS). Strain was elevated RAP (red arrow).

S1 Video. Movie of jaw movement at 5dpf control larvae expressing *Col2a1:mcherry.*

S2 Video. Movie of jaw movement at 5dpf anaesthetised larvae expressing *Col2a1:mcherry*.

S3 Video. Movie of jaw movement at 5dpf in double ablated larvae expressing *Col2a 1:mcherry*

